# Spatial modeling of tidal and land use impacts on *Escherichia coli* contamination and fluctuation in Asia’s urban river environmental gradient

**DOI:** 10.1101/2021.05.16.444347

**Authors:** Andri Wibowo

## Abstract

In the river, pathogens are the leading cause for rivers to exceed water and health quality standards. The GIS based spatial modeling and analysis were conducted to estimate *Escherichia coli* (*E. coli*) contaminated river bodies based on environmental and spatial gradients such as dissolved oxygen, pH, tidal, temperature and current in Asia’s urban river located in Kapuas river of Kalimantan. The *E. coli* was sampled from the river mouth up to the upstream land uses dominated by residential. The *E. coli* contamination was higher in the river mouth and in residential area as well. Likewise, *E. coli* contamination was higher during the low tide than high tide. During the high tide, the *E. coli* contamination were significantly affected by temperature and current (r^2^>0.5). Meanwhile, during the low tide, there were no dominant environmental factors that affect *E. coli* contamination. Hence, by knowing the spatial model of *E. coli* contamination driven by tidal, land use and environmental gradients, this paper has contributed to the advance management of water and river system.

## 1. Introduction

The major causes of water pollution is the presence of domestic waste in the waters [1]. The most dangerous part of domestic waste is pathogenic microorganisms contained in feces because it can transmit various diseases to human. The number of pathogenic microorganisms in the water is affected by tidal [2]. Correspondingly, pathogenic microorganism number is high especially in the low tide. During the low tide, the volume of water is low. Hence the ability to dilute water containing pathogenic microorganisms is decreasing. In contrast, during the high tide, the volume of water is high and the ability to dilute water containing pathogenic microorganisms is increasing. As a result, the pathogenic microorganism number is low.

Currently, the increasing activity of the population along the Kapuas river, an Asia’s urban river in Kalimantan, such as the increase in population settlements, the existence of markets and others, which generally dispose waste into the waters has affected river water quality. Correspondingly, there are high content of microorganisms detected in Kapuas river such as *Escherichia coli* (*E. coli*) and *Enterobacter aerogenes* [3]. The *E. coli* colony in Kapuas river was detected ranging from 14 MPN/100 ml to 1100 MPN/100 ml.

Geographic Information System (GIS) based spatial modeling has been viewed as solution to study the dynamic of *Escherichia coli* (*E. coli*) in the environment, especially in the water bodies. In Brazil [4], GIS methods were successfully applied to create spatial datasets for logistic regression model building and to construct risk maps using regression coefficients. This spatial model was used to study water quality problems and risk of waterborne enteric diseases in a lower-middle-class urban district. According to Kyna et al. [5], the highest potential *E. coli* loads were in the areas closest to the residential and the highest transported loads were located in the downstream.

Kapuas river is an Asia’s urban river located in West Kalimantan province and near the sea and its river mouth or estuary is bordered with Karimata strait and South China sea and hence affected by tides. Moreover, the land use near Kapuas river currently was occupied by the dominance of human settlements. The combination of tides and land use may affect significantly the *E. coli* in the river. Hence, this research aims to model spatially the effect of both tides and land use on the *E. coli* contamination in the Kapuas river.

## 2. Methodology

The data in this research were collected directly from the sampling stations and continued with analysis in the laboratory, especially for the *E. coli* samples. There were 10 sampling stations (Figure 1.a) located from the upstream to downstream closed to estuary of Kapuas river and sea. The stations were located from station a in upstream (latitude: −0.094316, longitude: 109.40594) to station j (0.054917, 109.172541) in downstream near estuary bordered with sea.

**Figure 1.a.**
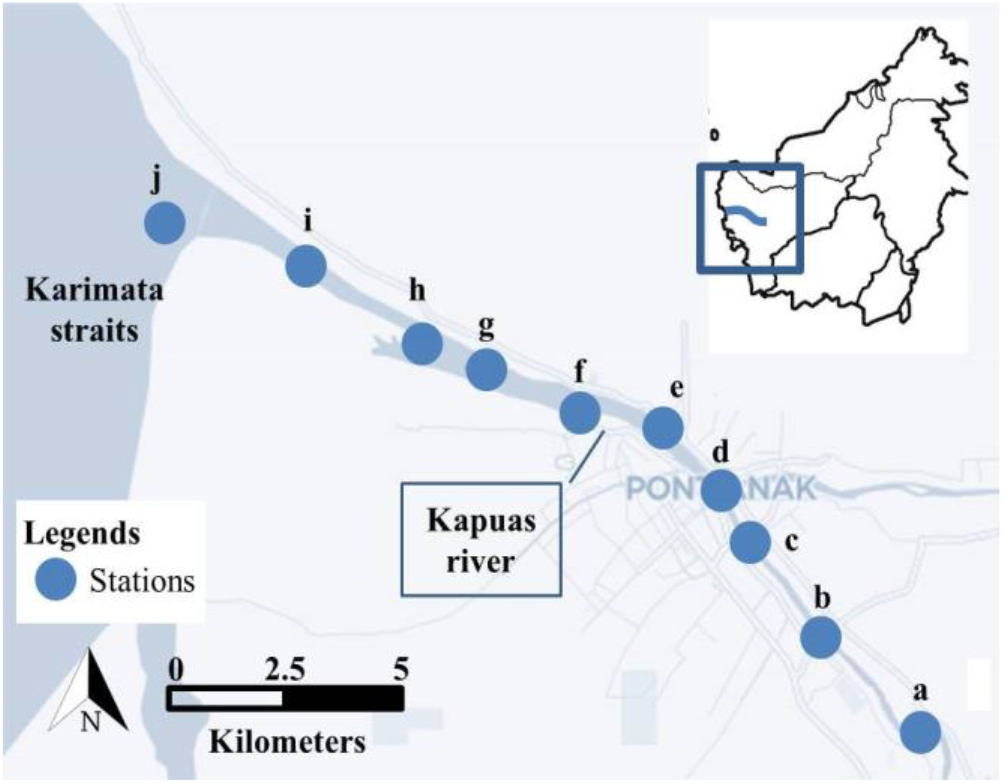
The locations of 10 sampling stations across Kapuas river.

### 2.1. Direct Measurement

The data that measured and collected directly were tides, dissolved oxygen (DO), pH, temperature, current, salinity and *E. coli*. The data collection methodology was following SNI 06-2412-1991 [6]. Daily tide data were recorded by using tide shaft that stationed near the station j located in the estuary. The *E. coli* samples were collected from the river by using 100 ml sterilized glass bottle. Simultaneously, the dissolved oxygen (DO), pH, temperature and current were measured. DO, pH and temperature of river water were measured by using multi parameter tools.

### 2.2. E. coli MPN Counting

The *E. coli* MPN (Most Probable Number) counting methodology was following SNI 6774: 2008 [7]. The 10 ml, 1 ml and 0.1 ml of river water each was inoculated in the tubes contained lactose broth. The tubes contained lactose broth and water then incubated at 37^0^c for 24 hours. After incubation, we counted how many tubes that show gaseous and color changes that indicate the *E. coli* presence. Then we converted the number of tubes containing *E. coli* with reference to MPN table to obtain the MPN value. To confirm and ensure the presence of *E. coli*, a sample from tubes then cultured in nutrient agar medium and incubated for 24 hours.

### 2.3 Spatial Mapping

All the sampling stations were georeferenced by using GPS to capture the coordinate of sampling stations. A GIS database table was developed with coordinate of sampling stations. This table has environmental parameter attributes including tides, dissolved oxygen (DO), pH, temperature, current, salinity and *E. coli.* By using GIS software, the environmental parameter layers were overlayed with the land use and Kapuas river layers. The *E. coli* layer was modeled by using interpolation method [8].

### 2.4. Salinity Gradient

The salinity was measured vertically from 0 m to 4 m depth with 1 m interval and horizontally from station j at the downstream estuary to station a at the upstream. The salinity data then were interpolated to generate the salinity depth gradient.

### 2.5. Statistical Analysis

The correlations of tides, dissolved oxygen, pH, temperature, current and *E. coli* were modeled by using correlation analysis.

## 3. Results and Discussion

The results show a gradient of land use across Kapuas river. In the downstream near estuary, the land use was dominated by dry land farming. In the middle part, the land use was dominated by settlements. Meanwhile, in the upstream, dry land with bushes dominated the land use (Figure 1.b). This gradient of land use has significantly affected the MPN of *E. coli* (Figure 2). The *E. coli* MPN was ranging from 31 x 10^3^ MPN/100 ml to 85 x 10^3^MPN/100 ml. The MPN was lower in the Kapuas river that surrounded with dry land farming in the downstream near estuary (station g, h, i, j) and upstream areas (station a, b, c). Meanwhile, the MPN of *E. coli* was significantly higher in Kapuas river that surrounded by settlements in midstream (station d, e, f). The finding in this research was in agreement with result from other researches. In Serin river, Sarawak, the anthropogenic activities have contributed to the *E. coli* concentrations in river with value ranged from 2 to 6.9 x 10^3^MPN/100 ml [9].

**Figure 1.b.**
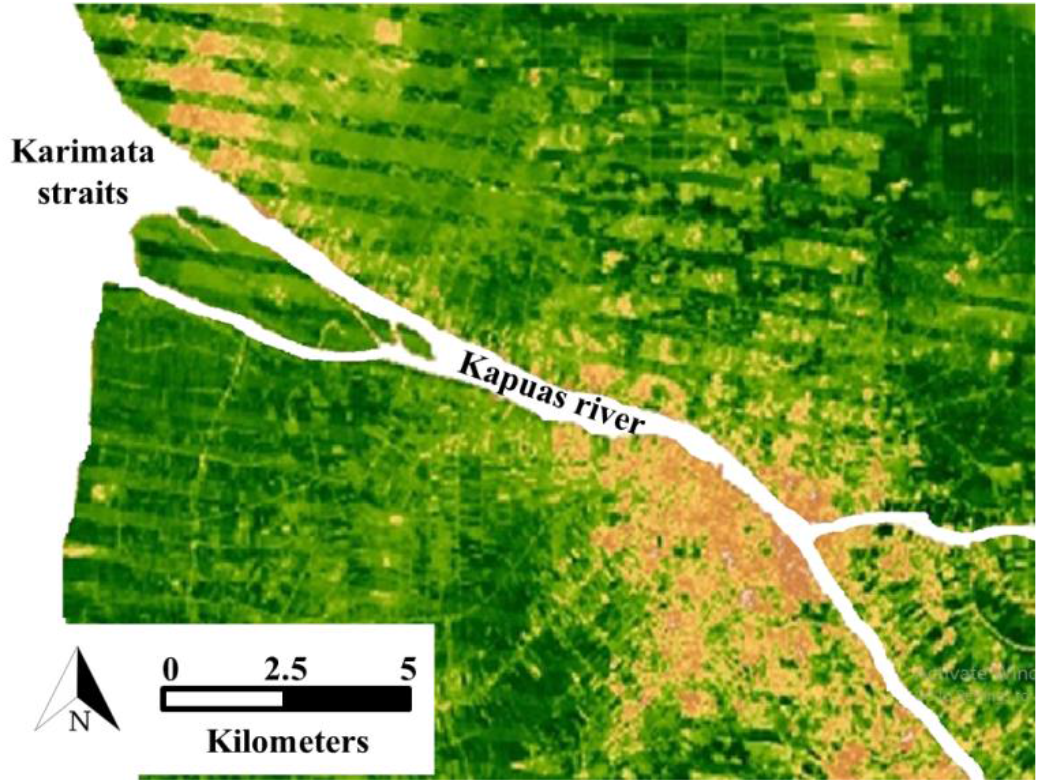
The land cover/use around 10 sampling stations in Kapuas river.

**Figure 2.**
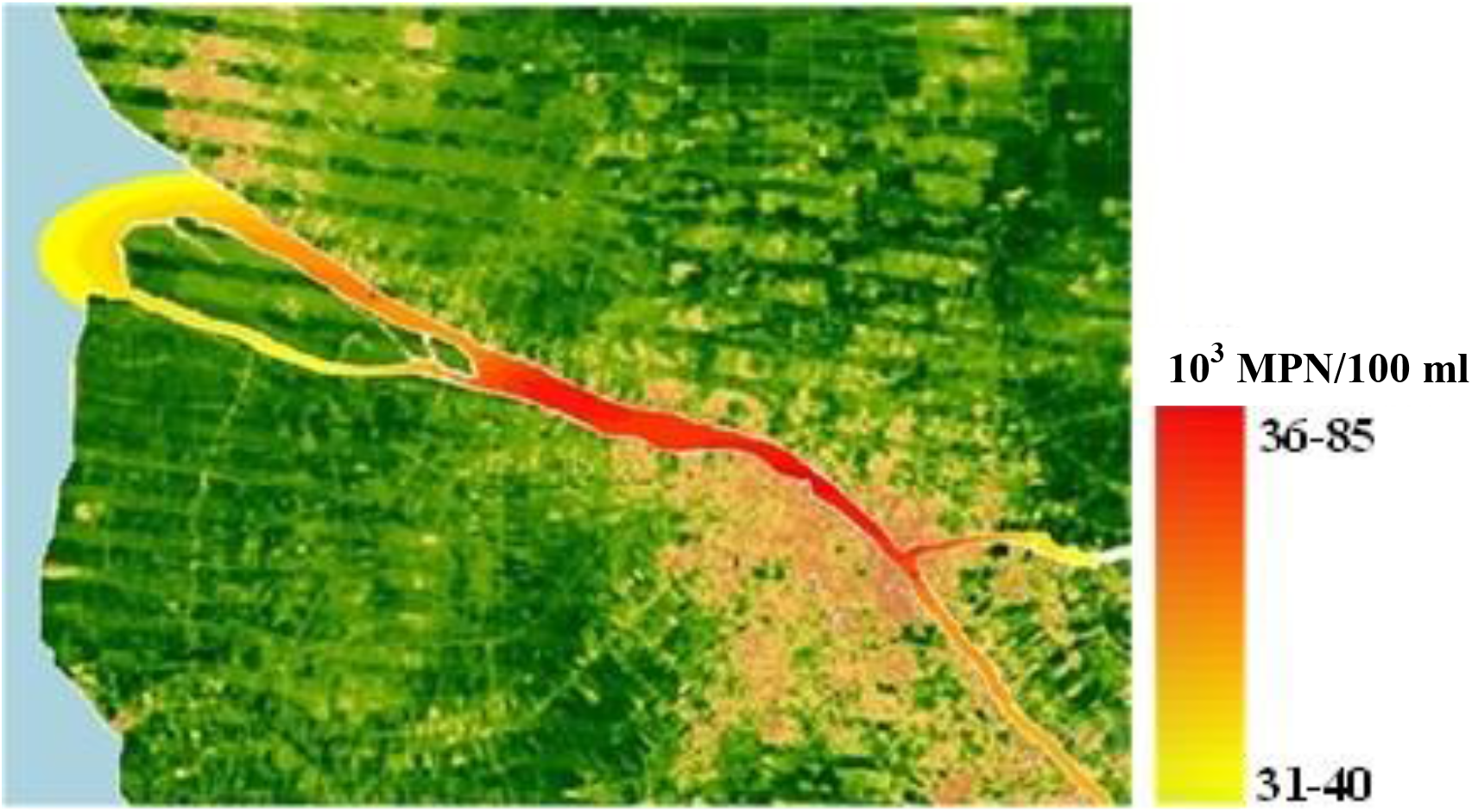
The model of *E. coli* MPN (10^3^ MPN/100 ml) gradients in 10 sampling stations across Kapuas river.

The salinity of Kapuas river was affected by the distance to the sea and depth as well. The salinity was higher in the downstream estuary at station j and the salinity was more higher than at the 4 m depth (Figure 3). At the surface, the salinity was ranging from 0 to 4 ppt. Meanwhile at 4 m depth, the salinity was ranging from 5 to 10 ppt. The salinity was also affected by the tide (p<0.05) at 4 m depth. Salinity was higher during the high tide compared to the low tide (Figure 3). During high tide, the sea water transported water that has high salt content and caused the water to be more saline.

**Figure 3.**
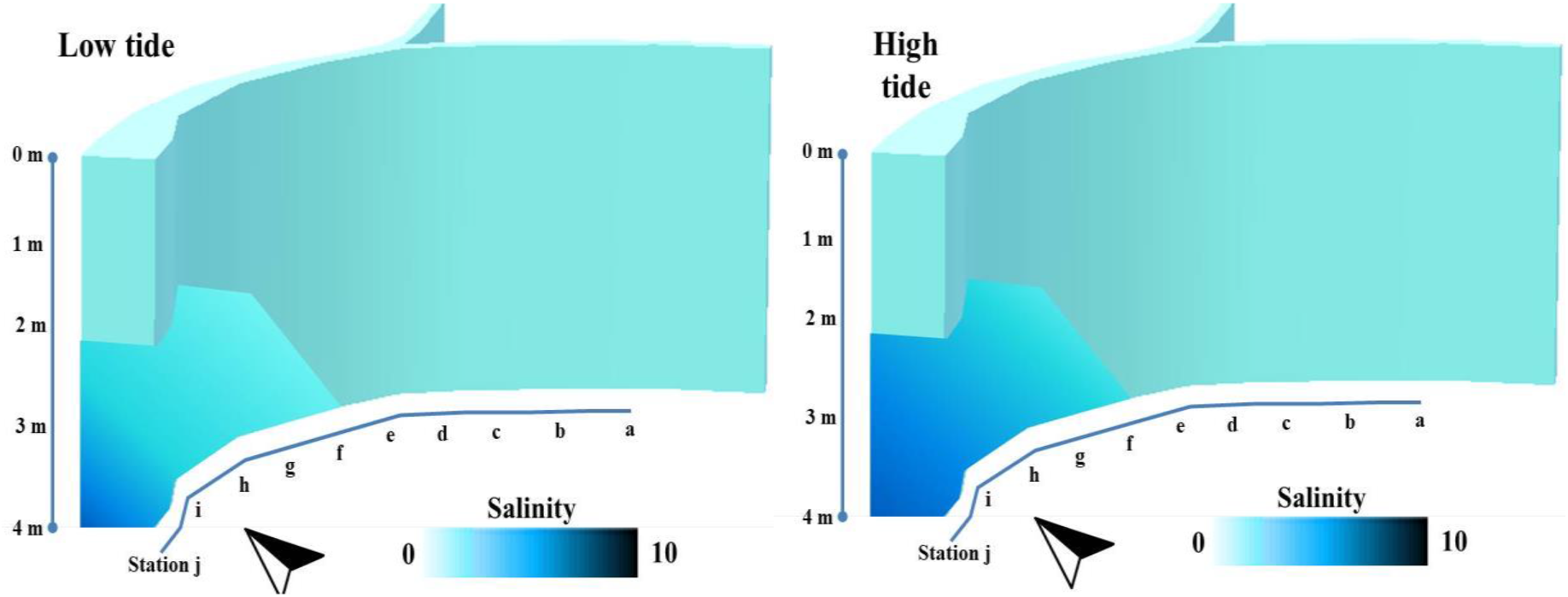
The isometric view of Kapuas river. This view depicts salinity (ppt) gradient model during low tide and high tide from 0 to 4 m depth at each station (a-j).

The *E. coli* in Kapuas river was also influenced significantly by the tide (Figure 4). The *E. coli* MPN was higher during the low tide than high tide. During the low tide, the salinity was low because volume of sea water containing salt was decreasing (Figure 3). The low salinity environment was preferable for *E. coli* to adapt. In water, *E. coli* prefers fresh water and has tolerance limit to salinity which is up to 37 ppt (10). Hence, during the low tide when the river contains more fresh water, *E. coli* MPN was higher. Respectively, during the high tide when the river contains more salt water and salinity was increasing, *E. coli* MPN was lower.

**Figure 4.**
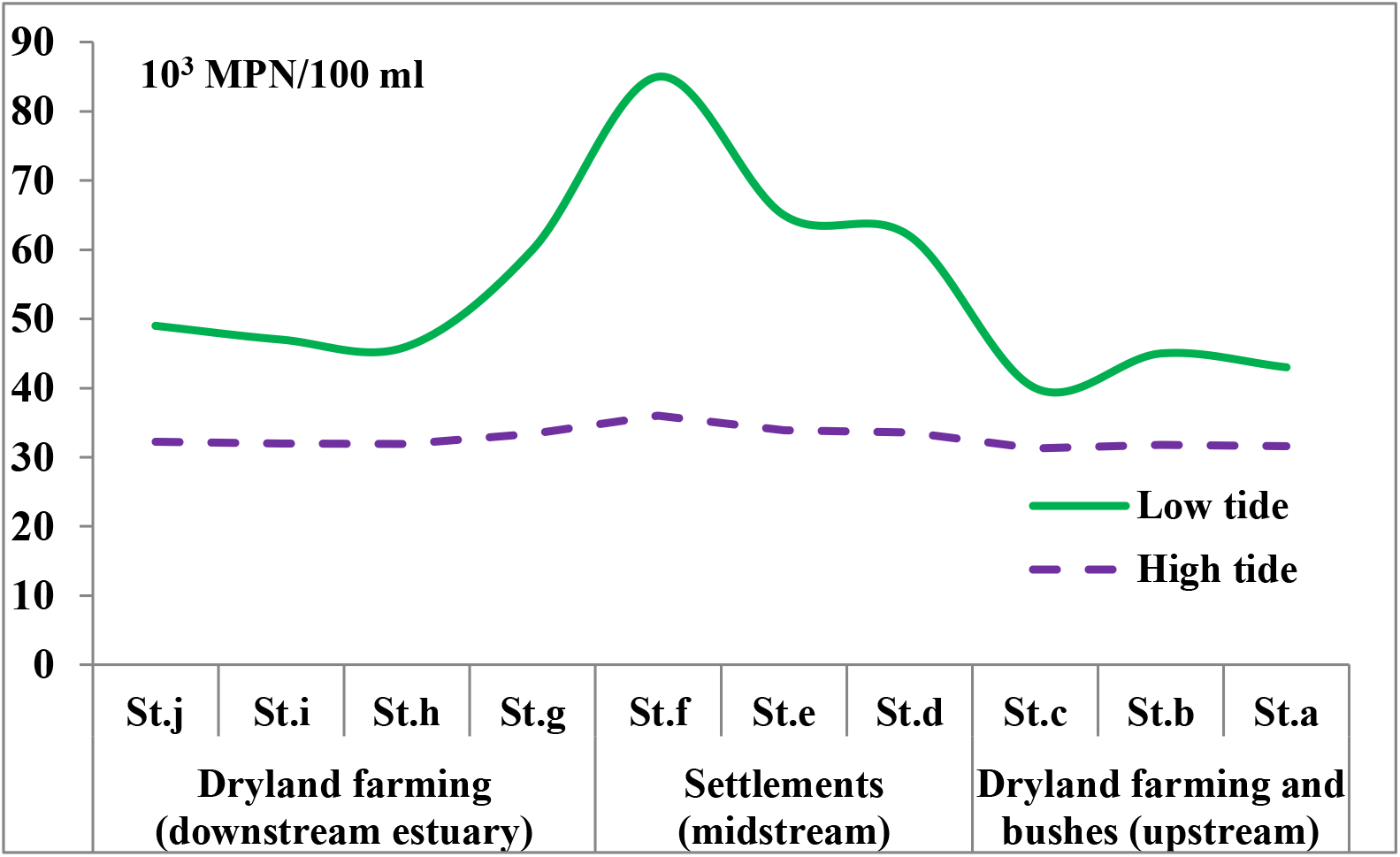
The effect of tide and land use on *E. coli* MPN in 10 sampling stations across Kapuas river.

During the low tide, there was no single factor that significantly affects the *E. coli* MPN. However, during the high tide, the *E. coli* MPN was affected significantly by temperature and current (Table 1). In high tide, increase in temperature will increase the *E. coli* MPN in river. Conversely, the increase in current will decrease the *E. coli* MPN.

**Table 1.** Correlation of MPN with temperature, DO, pH and current during low and high tide

|  | Temp. | DO | pH | Current |
| --- | --- | --- | --- | --- |
| MPN in low tide | 0.2591 | 0.0006 | -0.3897 | 0.3782 |
| MPN in high tide | 0.9929* | 0.3539 | 0.0098 | -0.9055* |

\*: significant correlation

## 4. Conclusions

The *E. coli* in Kapuas river was influenced by land use and tides as well. In settlements, *E. coli* was higher than in other places. *E. coli* MPN was also higher during low tide. The low tide provides more fresh water than salt water and reduces salinity, hence the river water is more suitable for *E. coli*.

Temperature and current are environmental parameters that can affect *E. coli*. Hence, by understanding and controlling these parameters, it can mitigate the potential of *E. coli* contamination in future.

